# Human lateral Frontal Pole contributes to control over social-emotional action

**DOI:** 10.1101/584896

**Authors:** Bob Bramson, Davide Folloni, Lennart Verhagen, Bart Hartogsveld, Rogier B. Mars, Ivan Toni, Karin Roelofs

## Abstract

Regulation of emotional behavior is essential for human social interactions. Recent work has exposed its cognitive complexity, as well as its unexpected reliance on portions of the anterior prefrontal cortex (aPFC) also involved in exploration, relational reasoning, and counterfactual choice, rather than on dorsolateral and medial prefrontal areas involved in several forms of cognitive control. This study anatomically qualifies the contribution of aPFC territories to the regulation of social-emotional actions, and explores a possible structural pathway through which emotional regulation might be implemented.

We provide converging evidence from task-based fMRI, diffusion-weighted imaging, and functional connectivity fingerprints for a novel neural element in emotional regulation. Task-based fMRI in human male participants (N = 40) performing a social-emotional approach-avoidance task identified aPFC territories involved in the regulation of social-emotional actions. Connectivity fingerprints, based on diffusion-weighted imaging and resting-state connectivity, localized those task-defined frontal regions to the lateral frontal pole (FPl), an anatomically-defined portion of the aPFC that lacks a homologous counterpart in macaque brains. Probabilistic tractography indicated that 10-20% of inter-individual variation in social-emotional regulation abilities is accounted for by the strength of structural connectivity between FPl and amygdala. Evidence from an independent replication sample (N = 50; 10 females) further substantiated this result. These findings provide novel neuroanatomical evidence for incorporating FPl in models of control over human social-emotional behavior.

**Significance statement:** Successful regulation of emotional behaviors is a prerequisite for successful participation in human society, as is evidenced by the social isolation and loss of occupational opportunities often encountered by people suffering from emotion-regulation disorders such as social-anxiety disorder and psychopathy. Knowledge about the precise cortical regions and connections supporting this control is crucial for understanding both the nature of computations needed to successfully traverse the space of possible actions in social situations, and the potential interventions that might result in efficient treatment of social-emotional disorders. This study provides evidence for a precise cortical region (FPl) and a structural pathway (the ventral amygdalofugal bundle) through which a cognitively complex form of emotional action regulation might be implemented in the human brain.

## Introduction

Control of social-emotional behavior is paramount for successful participation in society (Hare, 2017). However, regulating social-emotional actions is cognitively complex. It requires the ability to assess the effectiveness of on-going behavior and compare this with alternative action strategies. For instance, a scientist presenting her work in front of a critical audience needs to overcome the tendency to avoid a potential criticism in order to reap the long-term benefits of peer-exposure and constructive feedback. Recent converging evidence indicates that this type of regulation of social-emotional behavior is implemented by the rostral part of the anterior prefrontal cortex (aPFC; Volman et al. 2011, 2013; Koch et al. 2018). The current study anatomically qualifies the contribution of aPFC regions to the regulation of social-emotional action, and explores a structural pathway through which this regulation might be implemented.

The aPFC has been involved in a disparate range of cognitive tasks, e.g. relational reasoning (Vendetti and Bunge, 2014; Hartogsveld et al., 2017), counterfactual choice (Boorman et al., 2009a; Mansouri et al., 2017), exploration behavior (Daw et al., 2006; Zajkowski et al., 2017). To date, emotional action regulation has not been considered part of the cognitive fingerprint of this region (Koch et al., 2018). Models of emotion regulation have predominantly focused on dorsolateral and medial prefrontal areas (Etkin et al., 2015; Morawetz et al., 2017; Langner et al., 2018). These models are largely based on explicit regulation strategies (e.g. reappraisal; Wager et al., 2008; Buhle et al., 2014) and have ignored control of emotional actions, an important component of emotion regulation concerned with conflict between the emotional value of stimulus and response (Frijda et al., 2014; Ridderinkhof, 2017; Bramson et al., 2018). At first glance, the focus of the emotion control literature on medial prefrontal cortex (mPFC) is justified by the strong structural and functional connectivity between mPFC and areas involved in emotional and social processing, such as the amygdala (Ghashghaei and Barbas 2002; Petrides and Pandya 2007; Neubert et al. 2014; Tillman et al. 2018). In contrast, structural connections to the amygdala are scarce for anterior prefrontal areas (Ghashghaei et al., 2007). However, recent studies have shown that the amygdala is connected to the aPFC via rostral projections of the ventral amygdalofugal pathway (Krüger et al., 2015; Kamali et al., 2016). In fact, there is evidence that this tract might extend to the lateral frontal pole (FPl; Folloni et al., 2019), a portion of the human aPFC that does not have a homological counterpart in macaque brains (Neubert et al., 2014). These anatomical observations, together with converging functional evidence on aPFC involvement in the regulation of emotional strategies and actions (Bramson et al., 2018; Koch et al., 2018) raise the possibility that human social-emotional control is coordinated by the FPl.

In this study, we combine evidence from task-based fMRI, diffusion-weighted imaging, and resting-state connectivity fingerprints to define which portion of aPFC supports social-emotional control. First, we characterize the functional specificity of aPFC contributions to social-emotional control through a social-emotional approach-avoidance (AA) task. Participants approached or avoided happy and angry faces by means of speeded joystick reactions. Approaching angry and avoiding happy faces requires control over habitual social-emotional action tendencies, and elicits activity in aPFC (Roelofs et al., 2008; Volman et al., 2011b). The social relevance of the emotional-control indexed by this task has been established in social-emotional disorders (Bertsch et al., 2017; Volman 2016) and in healthy individuals where it predicted responses to real-life social stress-induction (Kaldewaij et al., 2019a). Second, we anatomically localize aPFC territories activated by the approach-avoidance task by matching structural and functional connectivity fingerprints of this area to fingerprints extracted from known areas of the aPFC (Neubert et al. 2014). Third, we assess the evidence for and the functional relevance of a structural pathway through which the portion of the aPFC activated by the AA task might support social-emotional regulation, namely through direct connections with the amygdala (Folloni et al., 2019).

## Methods

### Participants

Forty male students from the Radboud University Nijmegen participated in this study after giving informed consent. The sample had a mean age of 23.5 years (sd: 2.8, range 18-33 years). None of the participants reported history of mental illness or use of psychoactive/corticosteroid medication. All had normal or corrected to normal vision, and were screened for counter indications for magnetic resonance imaging (MRI). The study was approved by the local ethics committee (CMO:2014/288). Sample size was based on behavioral congruency effects observed previously in the same task (effect size *d = .4*; Volman et al, 2011). The sample consisted of males only in order to minimize potential variability caused by sex differences and fluctuations in cortisol and testosterone (e.g. due to different phases in the menstrual cycle). These hormones are known to influence behavioral and neural responses on the AA task (van Peer et al., 2007; Volman et al., 2011b; Kaldewaij et al., 2019a) and controlling for those fluctuations would have required a larger sample.

Given the novelty of one of the main findings of this report (correlation between amygdalofugal-FPl connectivity and behavioral congruency effect – see Results), we set out to replicate that finding in an independent convenience sample consisting of 50 participants [10 females, mean age = 23.8 (sd = 3.4, range 18-34)]. These participants took part in another study where they performed the AA task in a similar test-context, as well as a Diffusion weighted imaging (DWI) session. This second study was performed as part of a larger research project, after the analyses of the current study were finalized.

### Materials and apparatus

Images were acquired using a 3T MAGNETROM Prisma MRI scanner (Siemens AG, Healthcare Sector, Erlangen, Germany) using a 32 channel headcoil for functional and DWI images and a 20 channel headcoil for structural T1 images. Stimuli were presented using an EIKI LC-XL100 beamer with a resolution of 1024 × 768 and a refresh rate of 60 Hz, and were projected onto a screen behind the scanner bore. Participants were able to see the screen via a mirror.

The approach-avoidance (AA) task consisted of 16 blocks of 12 trials each, in which participants had to approach or avoid equiluminant happy or angry faces presented in the centre of the screen. Faces were presented for 100 ms and preceded by a 500 ms fixation cross in the centre of the screen. After the presentation of the face, participants had 2000 ms to respond using a joystick. Each trial was followed by an intertrial interval of 2 – 4 seconds. Task instructions for congruent (move the joystick towards you/away from you for happy/angry faces), and incongruent blocks (move the joystick towards you/away from you for angry/happy faces), were presented before each block and switched between blocks. Subjects responded using a customized fiber optic response joystick, which was fixed to only move in the sagittal plane. Affect-incongruent trials have consistently been shown to activate the aPFC (Roelofs et al. 2008; Volman et al. 2011; Tyborowska et al. 2016; Kaldewaij et al., 2019). Behavioral and neural responses on this task are affected in several social-emotional disorders such as psychopathy, social-anxiety and borderline disorder (Heuer et al., 2007; von Borries et al., 2012; Roelofs and Cremers, 2015; Bertsch et al., 2018), and those responses are influenced by hormones affecting social behavior (Volman et al., 2016; Kaldewaij et al., 2019a).

### Procedure

Data were acquired on two different days; day one consisted of the AA task, resting-state and T1 structural scan. DWI was acquired on the second day. Before and after the AA task we took saliva measurements to possibly control for testosterone and cortisol for other research purposes. On the second day, each participant filled out the State Trait Anxiety Inventory to control for effects of trait anxiety.

### Functional scans

The field of view of the scans acquired in the two MR-sessions was aligned to a built-in brain-atlas using an auto-align head scout sequence. During resting state and task fMRI sessions we acquired BOLD-sensitive images using a multiband sequence with TR = 735 ms/TE = 39ms, 64 slices, flip angle of 52°, multiband acceleration factor of 8, slice orientation T > C, voxel size = 2.4 × 2.4 × 2.4 mm, phase encoding direction A >> P. The resting state session lasted 8.5 minutes (700 images). Each sequence was followed by a field map image; flip angle = 60°, TR = 614, TE = 4.92.

### Structural scan

High-resolution anatomical images were acquired with a single-shot MPRAGE sequence with an acceleration factor of 2 (GRAPPA method), a TR of 2400 ms, TE 2.13 ms. Effective voxel size was 1 × 1 × 1 mm with 176 sagittal slices, distance factor 50%, flip angle 8°, orientation A ≫ P, FoV 256 mm.

### DW imaging

Diffusion-weighted images were acquired using echo-planar imaging with an acceleration factor of 2 (GRAPPA method). We acquired 65 2 mm-thick axial slices, with a voxel size of 2 × 2 × 2 mm, phase encoding direction A >> P, FoV 220 mm. In addition, we acquired 10 volumes without diffusion weighting (b = 0 s/mm^2^), 30 isotropically distributed directions using a b-value of 750 s/mm^2^, and 60 isotropically distributed directions a b-value of 3000 s/mm^2^. We also acquired a volume without diffusion weighting with reverse phase encoding (P >> A).

Diffusion-weigthed images for the replication sample were acquired using echo-planar imaging with multiband acceleration factor of 2 (GRAPPA method), multiband acceleration factor = 3. For this dataset we acquired 93 1.6 mm thick transversal slices with voxel size of 1.6 × 1.6 × 1.6 mm, phase encoding direction A >> P, FoV 211 mm, TR = 3350, TE = 71.20. 256 isotropically distributed directions were acquired using a b-value of 2500 s/mm^2^. We also acquired a volume without diffusion weighting with reverse phase encoding (P >> A). Preprocessing and analyses for this dataset were the same as for the original sample (reported below).

### Behavioral analyses

Behavioral analyses of AA task performance contrasted incongruent to congruent trials, comparing reaction times and percentage correct between conditions. Differences between conditions were assessed using repeated measures ANOVA and paired-sample t-tests.

### Analyses - functional images

All processing of the AA and resting state images was done using MELODIC 3.00 as implemented in FSL 5.0.10 (https://fsl.fmrib.ox.ac.uk). Images were motion corrected using MCFLIRT (Jenkinson et al., 2002), and distortions in the magnetic field were corrected using fieldmap correction in FUGUE. Functional images were rigid-body registered to the brain extracted structural image using FLIRT. Registration to MNI 2mm standard space was done using the nonlinear registration tool FNIRT. Images were spatially smoothed using a Gaussian 5 mm kernel and low passed filtered with a cut-off at 100 s. Independent component analysis was run (Beckmann and Smith, 2004), after which the first 10 components were manually inspected to remove sources of noise (Griffanti et al., 2017).

First and second level analyses were done in FEAT 6.00 implemented in FSL 5.0.10. We set up separate general linear models for each task. The first-level AA model consisted of four task regressors: Approach angry, approach happy, avoid angry and avoid happy; six motion regressors and their temporal derivatives; and two regressors modeling fluctuations in signal in white matter and cerebrospinal fluid. Approach happy and avoid angry (congruent conditions) were then contrasted with approach angry and avoid happy (incongruent conditions). This contrast was taken as input for each subject in the second level analysis. Second level analysis was performed using FMRIB’s Local Analysis of Mixed Effects (FLAME 1) with outlier de-weighting. We used a cluster forming threshold of Z > 2.3, a value that has been shown to effectively control for Family Wise Error rate, possibly related to the fact that FLAME estimates and takes into account within subject variance (Eklund et al., 2015). Locations with above threshold activity were localized using masks for the prefrontal cortex (Sallet et al., 2013; Neubert et al., 2014), parietal cortex (Mars et al., 2011) and whole brain cortical and subcortical atlases (Harvard-Oxford atlas).

To assess psychophysiological interactions (PPI) during the AA task we ran a second analysis using FEAT with regressors representing the task effects of interest (incongruent-congruent), the regressor incongruent + congruent, a regressor describing the first eigenvariate time series of activation of the seed region: Bilateral amygdalae taken from the AAL atlas (Tzourio-Mazoyer et al., 2002), and a regressor describing the interaction between the seed region activity and task effects. Strength of amygdalofugal-FPl connections as estimated from tractography of dw-MRI (see below) was added as a covariate in the second level analysis.

### Analyses – functional connectivity fingerprints of FPl based on resting-state fMRI

For the resting state fMRI analysis we defined regions-of-interest for each of the three areas in aPFC (FPl, FPm, and 46), as identified by Neubert et al. (2014). These masks were thresholded to contain only the 25% of voxels that were most likely to be part of each area (Mars et al., 2016; Hartogsveld et al., 2017). The masks were warped to each individual anatomical space, after which we extracted the first eigenvariate time series from each area. These time series were correlated with whole brain resting state activity, the results of which were transformed to z-values using Fishers z-transform and normalized. Resulting correlation maps were transformed back to MNI space after which we extracted the connectivity from each prefrontal region with five downstream regions: Posterior Cingulate Cortex (PCC) [10 - 52 24], Intraparietal lobule (IPL) [48 - 46 48], temporal pole (TP) [34 12 - 36], ventromedial prefrontal cortex (vmPFC) [8 44 - 14] and dorsolateral prefrontal cortex (dlPFC) [46 20 40], for each hemisphere separately. Differential connectivity with these five areas has previously been shown to be appropriate to dissociate different cortical regions within aPFC (Mars et al. 2016; Hartogsveld et al. 2017). This procedure creates a “connectivity fingerprint” of each region, describing functional (and structural for DWI analyses) connections with other brain areas, and rests on the premise that each regions’ unique functional repertoire is determined by its connections with other regions (Passingham et al., 2002; Mars et al., 2018). We followed the same procedure for the two regions found in the aPFC during the AA task: 5 mm sphere around MNI [24 50 - 4] and MNI [-24 52 4]. The focus of this study on aPFC responses to congruency demands during performance of the AA task is related to a large body of work that has characterized and replicated those aPFC responses across numerous studies (Roelofs et al., 2008; Volman et al., 2011a; Tyborowska et al., 2016). That work has shown that activity in aPFC during incongruent trials is correlated with and necessary for successfully overriding automatic action tendencies (Volman et al., 2011a); is influenced by several neuromodulators such as testosterone and cortisol (van Peer et al., 2007; Volman et al., 2011b, 2016; Kaldewaij et al., 2019a); is augmented in social psychopathology (von Borries et al., 2012; Radke et al., 2013; Bertsch et al., 2017, 2018); and predicts real-world stress coping (Kaldewaij et al., 2019b). The added value and validity of the structural explanation provided in the current study depends on using the same functional comparison implemented in those previous studies, namely a direct contrast between incongruent and congruent conditions.

For the statistical analysis, we computed the “city block” or “Manhattan” distance between the connectivity profiles of the AA regions and the connectivity profiles of the three prefrontal masks, separately for each hemisphere. This measure consists of the sum of absolute distance between different connectivity profiles and has previously been used to describe the relative dissimilarity between different connectivity profiles (Sallet et al., 2013; Neubert et al., 2014; Mars et al., 2016). We compared these distance measures to a distribution of randomized distances that we created by permuting the region labels 10000 times, each time randomly swapping the target region labels (FPl; FPm; area 46) within participants and computing the Manhattan distance with the AA fingerprint. The result of this approach shows whether the connectivity profile of the aPFC regions differs from the connectivity profile of the region active during the AA task (Mars et al., 2016).

### Analyses – structural connectivity fingerprints of FPL based on diffusion-weighted MRI

All analyses of diffusion data were performed in FSL FDT 3.0 (https://fsl.fmrib.ox.ac.uk). We used TOPUP to estimate susceptibility artifacts using additional b = 0 volumes with reverse phase coding direction (Andersson et al., 2003). Next, we used EDDY (using the fieldmap estimated by TOPUP) to correct for distortions caused by eddy currents and subject movement (Andersson and Sotiropoulos, 2016). We used BedpostX to fit a crossing fiber model using default settings (Behrens et al., 2007).

Seed regions were defined for five major tracts by creating 5 mm spheres in the Cingulum Bundle (CB) MNI: [8 - 2 36], Superior Longitudinal Fasciculus 1 (SLF1), MNI: [16 - 2 54], SLF 2, MNI: [30 - 2 38], SLF 3, MNI: [44 - 2 26] and Uncinate Fasciculus, MNI: [32 −2 −10] and then warped to individual subject space. As with the functional connectivity described above, these specific fiber bundles were chosen because they have been shown to be differentially connected to distinct areas within the aPFC (Neubert et al., 2014; Mars et al., 2016; Hartogsveld et al., 2017). The procedure for Amygdalofugal connectivity is described below. For the comparisons in the left hemisphere the same coordinates were used with the x-axis coordinate inverted. ProbtrackX was run with default settings using these masks as seed, separately for each hemisphere using an exclusion mask at x = 0. The results from probtrackX were warped to MNI 1 mm space, log transformed, and normalized to the highest probability per tract to allow a direct comparison between tracts with all values ranging between 0 and 1. We computed connectivity profiles of FPl, FPm and area 46 by counting the number of times each tract ended in these regions. These connectivity profiles were compared to the connectivity profile of the region in the aPFC resulting from the AA task contrast (5 mm sphere at MNI: [24 50 −4] for right and MNI: [-24 52 4] for left). We computed Manhattan distance between the connectivity profiles of the AA task and the three prefrontal masks and permuted this 10,000 times, each time randomly swapping the target region labels (FPl; FPm; area 46) within participants.

### Analyses - Amygdalofugal connectivity

Connections between amygdala and aPFC were reconstructed using probtrackX. For this we used the procedure and masks provided by Folloni et al., (2019). We seeded these connections in the white matter punctuating the extended amygdala and substantia innominata: MNI: [−7 3 −9] and an all-coronal waypoint mask at y = 22. We used tractography masks to ensure the estimated seedlines extended only rostrally, at least up to the y = 25 coronal plane, and excluded CSF and between-hemisphere connections. Connection strength was normalized and log transformed within each participant. Next we extracted the total amount of times the tractography entered the FPl, FPm and area 46 regions-of-interest separately. These values were correlated with congruency effects on reaction time and percentage correct using Spearman’s correlation coefficient.

To exclude that the Amygdalofugal-FPl connections are influenced by (much stronger) projections of the Uncinate Fasciculus (UF), we reconstructed UF connectivity with FPl and correlated this with the behavioral congruency. To isolate the UF we placed a seed in the white matter rostro-laterally to the amygdala (MNI: [35 2 −22]), a coronal waypoint mask at y = 22 and exclusion masks in the CSF and between hemispheres. We also used partial correlation to regress out possible correlations influences of UF and state anxiety (STAI-Y2) in the amygdalofugal-behavioral correlations. Finally, we used an independent comparison sample to confirm the correlation between amygdalofugal-FPl connectivity and behavioral congruency.

## Results

The study had two main goals: i) localize aPFC responses evoked during control over social-emotional action tendencies in relation to an established anatomical parcellation of the aPFC; and ii) provide evidence for the functional relevance of amygdala-aPFC connectivity in implementing control over emotional-actions. We implemented a-priori determined analyses addressing these two goals, correcting the statistical inferences for multiple comparisons, either through cluster correction (fMRI task contrasts), or Bonferroni-correction (anatomical localization and tract-behavioral correlations). Furthermore, we implemented a number of follow-up exploratory analyses, prioritizing sensitivity over specificity, thus avoiding corrections for multiple comparisons. The exploratory nature of these analyses is marked in the text.

### AA-task - Behavioral costs of controlling social-emotional behavior

There were interaction effects between emotion and movement for reaction times; *F(1,39)* = 26.5, *p* < .001 and error rates; *F(1,39)* = 27.2, *p* < .001. Reaction times were longer for incongruent (M = 673 ms, SD = 103) than for congruent movements (M = 626 ms, SD = 128), t(39) = 5.12, *p* <.001 with an effect size of *d* = .4, CI [.22, 1.03], calculated as (M1-M2) / sd_pooled_]. Participants also made more errors during incongruent (M = 94.3 % correct, SD = 4.43) than congruent blocks (M = 97.1 % correct, SD = 2.96), *t*(39) = 5.21, *p* < .001; effect size *d* = .7, CI [.27, 1.2]. In addition to these interaction effects there were main effects of movement *F(1,39)* = 32.9, *p* < .001; and emotion *F(1,39)* = 7.7, *p* = .008 on reaction times, but not on error rates, both *p* > .15. Approach movements were faster (M = 626 ms, SD = 106) then avoid movements (M = 661 ms, SD = 111) and reactions to happy faces (M = 635, SD = 103) were faster than to angry faces (M = 652, SD = 114). Interaction effects are depicted in figure 1B. These effects illustrate the behavioral cost of applying cognitive control over prepotent social-emotional action tendencies.

**Figure 1:**
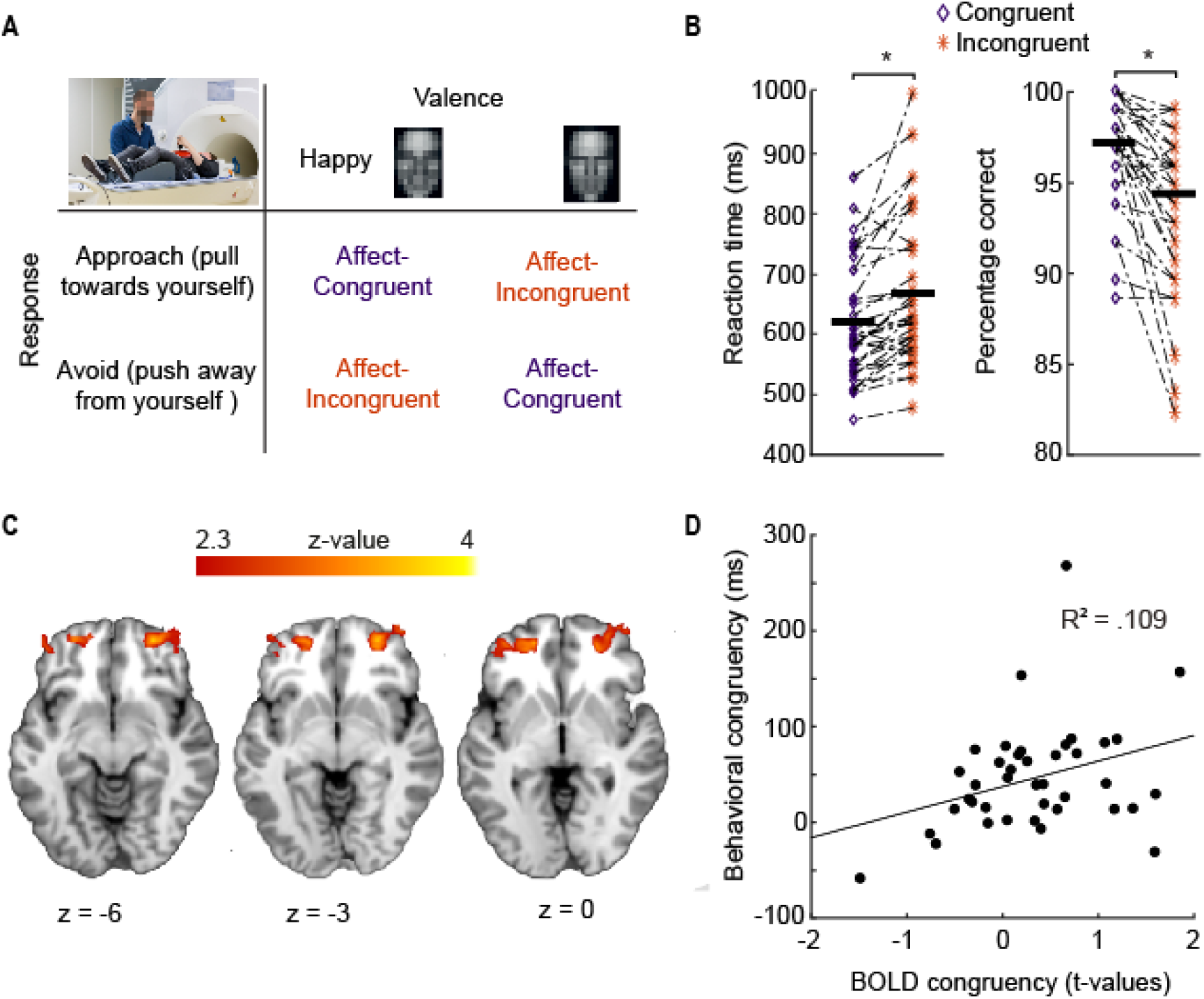
Effects of cognitive control over social-emotional action tendencies.

A) Schematic representation of the AA task used to measure control over social-emotional action tendencies. The incongruent condition requires control to override the automatic tendencies to approach happy- and avoid angry faces. B) Behavioral effects on AA task; participants show longer reaction times and more errors during the incongruent condition, illustrating the cost of exerting control. C) Main task effects for incongruent > congruent, masked to show above threshold (cluster level p < .001) activation in the anterior Prefrontal Cortex. (Frontal Pole mask; Harvard-oxford atlas) D) Correlation between reaction time congruency and BOLD congruency effects extracted from the lateral Frontal Pole across participants.

The replication sample also made more errors during incongruent (M= 94. 5 % correct) than in congruent trials (M = 96.8 % correct); *t*(49) = 6.25 *p* < .001.

### AA-task - Controlling social-emotional actions elicits frontal and parietal activation

Building on previous evidence (Volman et al. 2011; Tyborowska et al. 2016), we expected that control over social emotional action tendencies increases aPFC activity. This study tests whether this activity can be localized to the lateral Frontal Pole. We found clusters of increased activity over frontal areas during incongruent as compared to congruent trials, whole-brain cluster-level corrected for multiple comparisons (cluster-forming threshold of Z > 2.3 with corrected cluster threshold of *p* < .05). Overlaying the frontal activity profiles on the mask created by Neubert et al. (2014) confirmed activity in FPl, figure 1C. In addition we observed clusters of increased activity in bilateral intraparietal lobule, an area strongly connected to the aPFC (Mars et al., 2011); bilateral Insula/Inferior Frontal Gyrus [34 26 −6]; Bilateral area 8A [40 6 42]; bilateral area 46 [30 50 16]; and bilateral cerebellum [16 −78 −30].

Contrasting happy and angry showed increased activity for happy trials in cerebellum [4 −62 −50], posterior cingulate [8 −48 24], anterior cingulate [−2 40 2] and lateral occipital cortex [30 −84 0]. Avoid trials showed increased activity in Frontal orbital cortex [44 20 −10] and occipital cortex: [32 −66 54].

The estimated BOLD signal indexing control over emotional action tendencies (incongruent vs congruent trials) extracted from FPl correlated positively with the behavioral index of emotional control (reaction time differences between incongruent and congruent trials): *r(38)* = .33, *p* =.037, figure 1D. This effect corroborates our earlier findings of a positive relationship between individual differences in reaction time effects and recruitment of aPFC/FPl activation (Roelofs et al. 2009; Bramson et al., 2018). Exploratory analyses correlating BOLD congruency with behavioral congruency gave a trend correlation for area 46; *r(38)*= .31, *p* = .05 and no correlation for FPm; *r(38)* = .07, *p* = .63. These correlations did not differ from the congruency-FPl correlation, both *p* > .2.

### Relation between AA-related activity and functional fingerprints of FPl (Resting-state connectivity)

To assess whether the aPFC activity evoked during social-emotional control falls within FPl (Neubert et al. 2014), we compared resting-state connectivity profiles of area 46, FPl and FPm with the connectivity profile of the aPFC clusters showing a congruency effect (MNI: [24 50 −4] and [−24 52 −4]). We tested whether the connectivity fingerprints of the three prefrontal areas differed from the connectivity profile of the AA-related clusters, figure 2.

**Figure 2:**
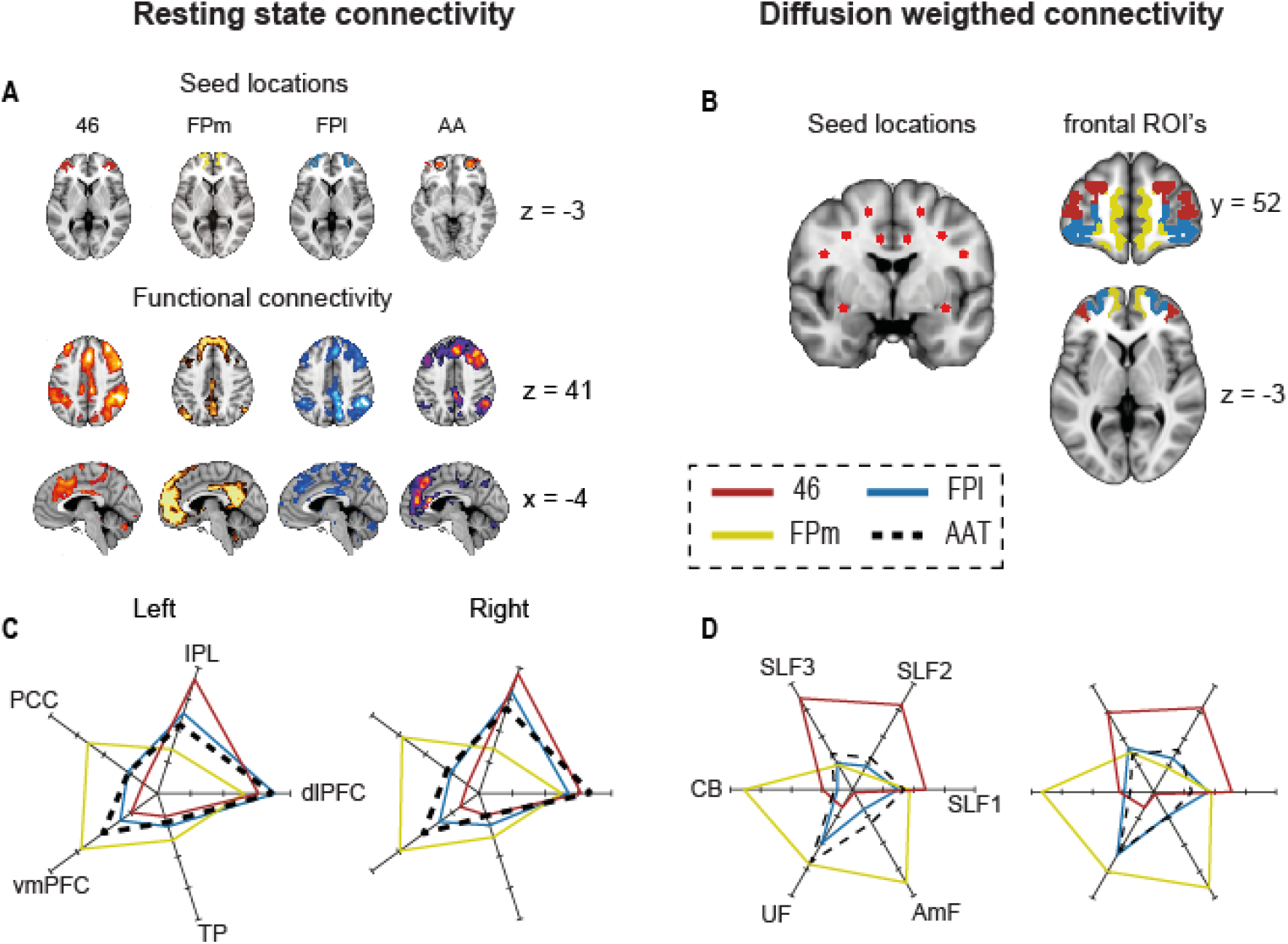
AA task effects are localized within the connectivity-defined lateral Frontal Pole.

A) Seed regions and whole brain resting-state connectivity for area 46, FPm, FPl, and for the cluster showing AA task-effects. B) Seed locations for tractography in five large white matter tracts (AmF seed not shown) and masks for the regions FPl, FPm and area 46. C) Resting state connectivity fingerprints of the aPFC and AA regions, separated for left and right ROI’s. Region labels: Posterior Cingulate Cortex (PCC), ventromedial prefrontal cortex (vmPFC), Intraparietal Lobule (IPL), dorsolateral Prefrontal Cortex (dlPFC), Temporal Pole (TP). D) Connectivity fingerprints for tractography of aPFC and the AA regions, separated for left and right ROI’s. Tract abbreviations: Superior Longitudinal Fasciculus (SLF1, SLF2, SLF3); Uncinate Fasciculus (UF); Cingulum Bundle (CB); Amygdalofugal (AmF).

Qualitative inspection of the resting-state fingerprints showed relatively similar connectivity profiles for area 46 and FPl. Both are strongly correlated with the intraparietal sulcus (IPl) and dorsolateral prefontal cortex (dlPFC); however, FPl shows stronger connectivity to the posterior cingulate cortex (PCC). Functional connectivity of FPm differed quite strongly from both area 46 and FPl, showing strongest connectivity with ventromedial prefrontal areas (vmPFC), PCC and temporal pole (TP), figure 2A;C. These results corroborate earlier reports on resting-state connectivity fingerprints for these areas (Neubert et al., 2014; Mars et al., 2016; Hartogsveld et al., 2017).

Comparing connectivity profiles between the three areas in aPFC with the connectivity profile extracted from the AA-related region showed that its fingerprint differed significantly from that of area 46, (left hemisphere: p < .001; right-hemisphere: p < .001), and from that of area FPm (right hemisphere: p < .001; left-hemisphere: p < .001), but not from the connectivity fingerprint of area FPl (left-hemisphere: p = .87, right-hemisphere: p = .075). This result provides evidence that control of social emotional action tendencies involves the lateral Frontal Pole.

### Relation between AA-related activity and structural fingerprints of FPl (DWI connectivity)

In addition to resting-state connectivity, we created connectivity profiles based on white matter tractography. We placed seeds in five large white matter tracts (figure 2B); performed probabilistic tractography to the rest of the hemisphere; and counted how often each tract ended in each of the three regions in aPFC (figure 2B). We used a similar procedure for amygdalofugal (AmF) connectivity (see methods).

Comparing connectivity profiles of the aPFC regions to these five major white matter tracts and AmF connectivity showed relatively strong connectivity between area 46 and the three superior longitudinal fasciculi (SLF1, SLF2 and SLF3; figure 2D). FPm is most strongly connected to the Cingulate Bundle, Uncinate Fasciculus and Amygdalofugal pathways. FPl is less strongly connected to the tracts leading outside the frontal cortex, and more strongly connected to the Uncinate Fasciculus, which leads to the temporal pole (among other regions). The AA-related regions are also more strongly connected to the Uncinate Fasciculus. These results are shown in figure 2D and match earlier reports on connectivity of aPFC regions (Neubert et al., 2014; Mars et al., 2016; Hartogsveld et al., 2017).

Comparing connectivity profiles between the three areas in aPFC with the connectivity profile extracted from the AA region showed that the fingerprint of the AA region differed significantly from that of area 46, (left hemisphere: p < .001; right-hemisphere: p < .001), and from that of area FPm (right hemisphere: p < .001; left-hemisphere: p < .001), but not from the connectivity fingerprint of area FPl (left-hemisphere: p = .98, right-hemisphere: p = 1). This result corroborates the resting state results, providing additional evidence that the FPl is one of the regions involved in controlling social emotional action tendencies.

### FPl-amygdalofugal connectivity partly explains performance on the AA task

To identify an anatomical pathway through which the FPl could support social-emotional behavior, we tested whether inter-subject variability in the strength of FPl structural connectivity mediated by the amygdalofugal connections predicts successful performance on the AA task. On average, the FPm is more strongly connected to amygdalofugal pathways than the FPl and area 46 (Figure 3A), in line with earlier studies showing both functionally and anatomically stronger connections between limbic areas and medial prefrontal regions (Ghashghaei and Barbas, 2002; Neubert et al., 2014; Folloni et al., 2019). However, individual variability in the connectivity of FPl and FPm confirms that the former region is more strongly involved in social-emotional control than the latter. Namely, there was a significant correlation between tract strength and AA congruency effect on percentage correct; Spearman’s *r(38)* .44 =, *p* = .004 for the FPl connections but not for FPm, *r(38)* = .24, *p* = .13 and area 46, *r(38)* = .3, *p* = .062, figure 3B. However, these correlations did not differ from one another, both comparisons *p* > .3. The direction of the FPl-behavioral correlation indicated that stronger connectivity was related to relatively worse performance when having to exert control. There were no significant correlations between tract strength and congruency effect on reaction time; all *p* > .09. The congruency-FPl correlation for percentage correct survives Bonferroni-correction for multiple comparisons over the eight correlations computed between behavioral congruency and amygdalofugal-FPl connectivity; and the two control correlations described below.

**Figure 3:**
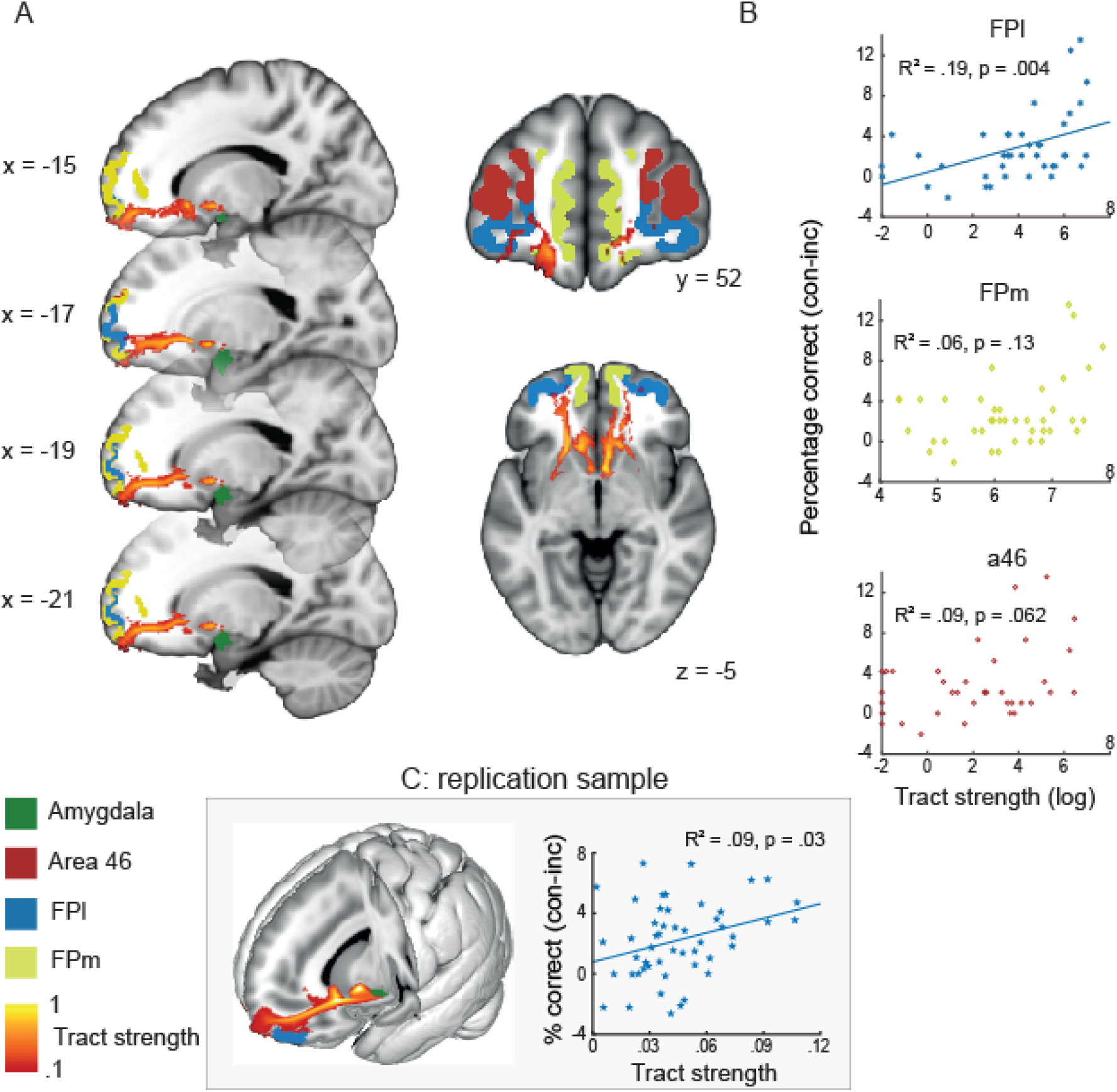
Amygdalofugal connections with FPl correlate with AA task performance.

A) Illustration of the reconstructed tract that leads from the Amygdala to the anterior Prefrontal cortex. This tract shows strongest connectivity with medial prefrontal areas, as can be expected given earlier studies. B) The number of times the tract ends in FPl correlates with behavioral congruency in error rates on the AA task, r = .44, p = .0044, suggesting that the amygdalofugal pathway is involved in mediating FPl-amygdala functional interactions during social-emotional control. C) Illustration of the reconstructed tract and correlation between behavioral congruency and tract strength with FPl in the replication sample.

To facilitate data interpretation, we conducted 2 additional exploratory control analyses. First, to exclude that the correlation between amygdalofugal-FPl connectivity and AA performance could be better explained by connections between amygdala and FPl through the Uncinate Fasciculus (UF; Petrides and Pandya 2007) or trait anxiety (Kim and Whalen, 2009), we correlated UF-FPl connectivity with AA performance, and amygdalofugal-FPl connectivity with scores on the State anxiety inventory (STAI). None of these analyses yielded significant correlations, all *p* > .58. Nor did the correlation between amygdalofugal-FPl connections and behavioral congruency change when controlling for state anxiety or UF-FPl connectivity.

Second, to explore whether FPl-amygdalofugal connectivity would also influence functional connectivity between these two areas, we performed a Psychophysiological Interaction (PPI) analysis with bilateral amygdalae as seed region and the strength of the connection with FPl as covariate. This analysis showed a negative correlation between tract strength and amygdala-FPl functional connectivity during the task, when using a small volume correction over bilateral FPl; *p* = .019 (FWE cluster corrected; cluster location MNI [−32 50 6], cluster size: 20 voxels), meaning that with increased tract strength there is more negative connectivity between the FPl and amygdala, which might be interpreted as increased regulation of amygdala by the FPl (Volman et al., 2013).

Finally, given the novelty of the role of the amygdalofugal path in emotion control, we tested whether the correlation between behavioral congruency and amygdalofugal connectivity would replicate in an independent sample of 50 participants. The positive relationship between the congruency effect in % correct on the AA task and amygdalofugal-FPl connections was confirmed; Spearman’s *r(48) = .3*, *p = .03*, figure 3C, further substantiating this finding.

## Discussion

The present study anatomically qualifies the involvement of the anterior prefrontal cortex in social-emotional action control. There are two main findings. First, structural and functional connectivity fingerprints of the lateral Frontal Pole (Neubert et al., 2014) closely match the fingerprints of the aPFC territory recruited when social-emotional control is required. Second, the strength of FPl structural connectivity with the amygdala accounts for a substantial portion of inter-individual variation in social-emotional regulation abilities. These findings provide evidence for a precise cortical region (FPl) and a structural pathway (the ventral amygdalofugal bundle) through which a cognitively complex form of emotional regulation might be implemented in the human brain.

## FPl supports social-emotional control

Previous work has repeatedly shown that the aPFC is involved in controlling social-emotional behavior, and that it is capable of doing so by influencing downstream activity in the amygdala (Volman et al. 2013; Tyborowska et al. 2016); motor cortex; and parietal areas (Volman et al. 2011; Bramson et al. 2018). Here, we extend this knowledge by showing that the aPFC contribution to social-emotional control arises from a precise anatomical structure: the lateral Frontal Pole (Neubert et al., 2014). This finding is based on the connectivity fingerprint of the FPl. This anatomical metric also provides clues to the information received by the FPl, and the influence it can exert on other brain structures (Mars et al., 2018). The FPl, differently from neighbouring areas FPm and 46, is connected to both medial and lateral circuits with parietal, cingulate, and temporal regions (Figure 2). The peculiar pattern of FPl connectivity fits with the known supramodal regulatory role of this area during social-emotional control (Volman et al. 2011; Volman et al. 2013; Koch et al. in prep), and with the notion that social-emotional regulation involves access to both egocentric value of emotionally-laden actions, as well as social consequences of those actions (Koch et al., 2018).

This study was designed to add functional specificity to those anatomically-grounded inferences on aPFC contributions to social-emotional regulations. The observations of this study link social-emotional regulation to several cognitive processes that rely on the FPl, e.g. concurrent monitoring of current and alternative goals (Burgess et al., 2007; Badre and D’esposito, 2009; Mansouri et al., 2017), cognitive exploration (Daw et al., 2006; Zajkowski et al., 2017), counterfactual reasoning (Boorman et al., 2009b; Koechlin, 2016), metacognition (Fleming et al., 2014; Shekhar and Rahnev, 2018), and relational reasoning (Vendetti and Bunge, 2014; Hartogsveld et al., 2017). More precisely, it becomes relevant to test whether the FPl contribution to social-emotional control is linked to the online maintenance and evaluation of alternative goals or counterfactual regulation strategies (Sheppes et al., 2014; Koch et al., 2018); or whether the cognitive demands of social-emotional control and their input-output connectivity segregate this faculty from other known FPl functions.

## Amygdala-FPl projections modulate control over emotional-action

This study shows that control over social-emotional actions is negatively associated with the strength of amygdala-FPl connections, with connections between amygdala and FPl explaining over 20% of the variance on the AA task. This result was confirmed in an independent replication sample. On the assumption that the amygdala-FPl connectivity indexed in this study captures the amygdalofugal pathway (Folloni et al. 2019), i.e. the main efferent pathways from amygdala to frontal cortex (Nolte, 1999; Ghashghaei and Barbas, 2002; Noback et al., 2005; Krüger et al., 2015; Kamali et al., 2016), this finding suggests that stronger amygdala projections to the FPl impair emotional regulation. Namely, during incongruent trials, stronger amygdala afferences to the FPl could lead to stronger automatic action tendencies or heightened emotional vigilance and thus more errors when those tendencies need to be rapidly over-ruled. This observation fits with previous imaging work showing that successful emotional control reduces bottom-up effective connectivity between amygdala and aPFC (Volman et al., 2013). Moreover, it suggests that top-down regulatory FPl efferences might reach the amygdala through other fiber bundles, e.g. the uncinate fasciculus (Petrides and Pandya, 2007; Folloni et al., 2019); or through other regions, e.g. the nucleus accumbens (Wager et al., 2008) or the orbitofrontal cortex (Ray and Zald, 2012).

The relation between amygdala-FPl connectivity strength and social-emotional action control observed in this study is surprisingly large when compared to the magnitude of previous structure-function correlations in the same anatomical regions (Jung et al., 2018). We suspect that the large effect size is a consequence of considering the anatomical differences between fiber bundles connecting the amygdala to the prefrontal cortex, and in particular the uniquely human configuration of that connectivity, since anthropoid monkeys do not possess a functional homologue of the human lateral Frontal Pole (Semendeferi et al., 2010; Neubert et al., 2014; Mars et al., 2016). Future tests of this potential functional-anatomical dissociation between bottom-up and top-down amygdala-FPl connectivity might also help to clarify the heterogeneity of previous findings concerning the relation between amygdala-prefrontal connectivity and state-anxiety measures (Kim and Whalen, 2009; Clewett et al., 2014).

## Interpretational issues

It could be argued that the regulation required in the AA task is just an instance of inhibitory control with social stimuli, and that regulation in this task could be accomplished by suppressing automatic emotional action tendencies (Etkin et al., 2006; Aron et al., 2014). Several studies have shown that this is not a viable option. First, when participants respond to the gender rather than the emotion of the face, both behavioral (congruency effect) and neural correlates of emotional control (FPl activation) are extinguished (Roelofs et al., 2008; Volman et al., 2011b). This illustrates that the regulation indexed by the AA task crucially depends on the interaction between emotional percepts and action selection, rather than on overriding conflict contained in the stimulus itself (e.g. emotional Stroop task, which is supported by different neural circuitry (Etkin et al., 2011, 2015)). Second, performance on the AA task is specifically altered in patients with disrupted social-emotional regulation (Heuer et al., 2007; van Peer et al., 2009; von Borries et al., 2012). For instance, aggression-related psychopathologies such as psychopathy are marked by reduction of avoidance responses, specific to angry faces, an effect related to measures of social aggression (von Borries et al., 2012). In contrast, patients with social anxiety disorder show a consistent increase of avoidance responses, specific to angry faces (Roelofs et al., 2005, 2009, 2010; Heuer et al., 2007). In addition, socially-relevant hormones such as testosterone (Tyborowska et al., 2016), cortisol (Volman et al., 2011b), and oxytocin (Radke et al., 2017) influence performance of the Approach-Avoidance task in healthy participants. Most critically, the aPFC congruency effect measured during the social approach-avoidance task is predictive of subjective, physiological and hormonal responses to a real-life social stressor (Kaldewaij et al., 2019b). In sum, these studies highlight the validity of the AA task for indexing social-emotional regulation.

In principle, the relation between strength of amygdala-FPl connectivity and behavioral emotional control could be explained by a third factor, e.g. trait anxiety. However, the tract strength between amygdala and FPl did not correlate with trait anxiety, and the correlation between behavioral congruency and tract strength persisted when controlling for anxiety. This observation confirms that, in healthy participants, aPFC activity during the AA task is orthogonal to trait anxiety effects (Bramson et al., 2018), in contrast to known trait anxiety effects on amygdala-vmPFC connectivity (Kim and Whalen, 2009).

It is not easy to acquire reliable fMRI signals from anterior prefrontal regions (Hutton et al., 2002). Therefore, we used magnetic field corrections to restore signal distortions around the frontal pole (Jezzard and Balaban, 1995; Jenkinson, 2004). The resting-state connectivity profiles closely matched those of earlier studies that used different MRI protocols (Neubert et al., 2014; Mars et al., 2016) arguing against the possibility that the results reported here can be attributed to signal distortions. The first sample of participants consisted of males only, and the replication sample was predominantly male. Whereas behavioral and neural effects in the AA task have been shown in mixed and female only samples as well (Radke et al., 2015; Tyborowska et al., 2016; Bertsch et al., 2018), we recruited only males to avoid having to control for differences in social hormones such as testosterone and cortisol, which would require a larger sample of participants. These hormones are known to influence behavioral and neural responses on the AA task (van Peer et al., 2007; Volman et al., 2011b; Kaldewaij et al., 2019a). Future studies could explore similarities and potential differences in emotional processing between sexes (Domes et al., 2010; Whittle et al., 2011).

## Conclusion

This study provides anatomical evidence supporting the involvement of the lateral frontal pole, partly via the ventral amygdalofugal pathway, in the regulation of social-emotional behavior (Koch et al., 2018). The findings have implications for structuring mechanism-based interventions in psychopathologies characterized by altered emotional control abilities, e.g. anxiety disorders and psychopathy (Volman et al., 2016). The findings are also relevant for understanding the neurobiological and cognitive complexities underlying rule-based regulation of action tendencies, arguably a crucial pre-requisite for the development of human cumulative culture (Hare, 2017; Whiten, 2017).

## Acknowledgements

This work was supported by a VICI grant (#453-12-001) from the Netherlands Organization for Scientific Research (NWO) and a consolidator grant from the European Research Council (ERC_CoG-2017_772337) awarded to Karin Roelofs. The work of RBM is supported by the Netherlands Organization for Scientific Research (NWO) [452-13-015]. The work of DF was supported by Wellcome Trust UK Grant (105238/Z/14/Z).

